# *In vivo* sequestration of innate small molecules to promote bone healing

**DOI:** 10.1101/745166

**Authors:** Yuze Zeng, Yu-Ru V. Shih, Gurpreet S. Baht, Shyni Varghese

## Abstract

Approaches that enable innate repair mechanisms hold great potential for tissue repair. Herein, we describe biomaterial-assisted sequestration of small molecules to localize pro-regenerative signaling at the injury site. Specifically, we designed a synthetic biomaterial containing boronate molecules to sequester adenosine, a small molecule ubiquitously present in the human body. The biomaterial-assisted sequestration of adenosine leverages the transient surge of extracellular adenosine following injury to prolong local adenosine signaling. We demonstrated that implantation of the biomaterial patch following injury establishes an *in-situ* stockpile of adenosine, resulting in accelerated healing by promoting both osteoblastogenesis and angiogenesis. The adenosine content within the patch recedes to the physiological level as the tissue regenerates. In addition to sequestering endogenous adenosine, the biomaterial is also able to deliver exogenous adenosine to the site of injury, offering a versatile solution to utilizing adenosine as a potential therapeutic for tissue repair.

## Introduction

A leading concept in regenerative medicine is transplantation of tissue-specific cells, often supported by biomaterials, to promote tissue repair^1^. While this strategy has achieved some success, its broad clinical application is hindered by various challenges such as high costs, constraints associated with cell isolation and expansion, and limited *in vivo* engraftment of transplanted cells^2-4^. Instead, mobilizing endogenous cells to augment the innate regenerative ability of tissues has been explored as an alternative^5-8^. Given that the function of endogenous cells is regulated by their microenvironment, the potential of biomaterials and/or growth factors to create pro-healing niches for endogenous cells has been explored extensively^6,9-14^. Meanwhile, naturally-occurring small molecules are also appealing and equally powerful in regulating various cellular functions including tissue-specific differentiation of stem cells^15-17^. Although significant strides have been made in employing small molecules to direct cellular functions *in vitro*, harnessing small molecules towards tissue repair *in vivo* still remains limited.

In this study, we determine whether sequestration of small molecules could be used to augment endogenous cell function leading to improved tissue repair. Adenosine is a small molecule ubiquitously present in the human body which acts as an extracellular signaling molecule through G-protein coupled adenosine receptors^18,19^. While the physiological concentration of extracellular adenosine is often insufficient to activate adenosine receptors^20^, an increase in extracellular adenosine is observed following tissue injury, which is integral to the natural repair mechanism^18,21-23^. However, this increase is transient as adenosine is rapidly metabolized^24,25^. Although delivery of adenosine can be employed to activate adenosine signaling and address tissue dysfunctions, in practice, such an approach has remained elusive. This is mainly due to the ubiquitous nature of adenosine and the potential off-target effects associated with its systemic administration^25-27^. Instead, approaches that localize adenosine signaling at the targeted tissue site can circumvent these limitations and open up new viable therapeutic strategies.

To this end, we have developed a biomaterial-based approach to sequester extracellular adenosine by capitalizing on its transient surge following trauma or by delivering exogenous adenosine to sustain the activation of adenosine signaling strictly at the injury site. Specifically, we leveraged the ability of boronate molecules to bind to adenosine via dynamic covalent bonding^28-30^ (**Fig. 1a**). By employing a 3-(acrylamido)phenylboronic acid (PBA)- functionalized polyethylene glycol (PEG) network, we demonstrated *in vivo* sequestration of adenosine and its application to accelerate bone repair in a murine model (**Fig. 1b**). We used bone fracture as a model, due to its clinical relevance^31-33^. Bone fracture is also well-suited for studying adenosine-mediated tissue repair, as extracellular adenosine and its receptors play a key role in maintaining bone homeostasis and function^20,22,34^ and have been proven to induce osteogenic differentiation of progenitor cells^17,35-37^.

**Figure 1.**
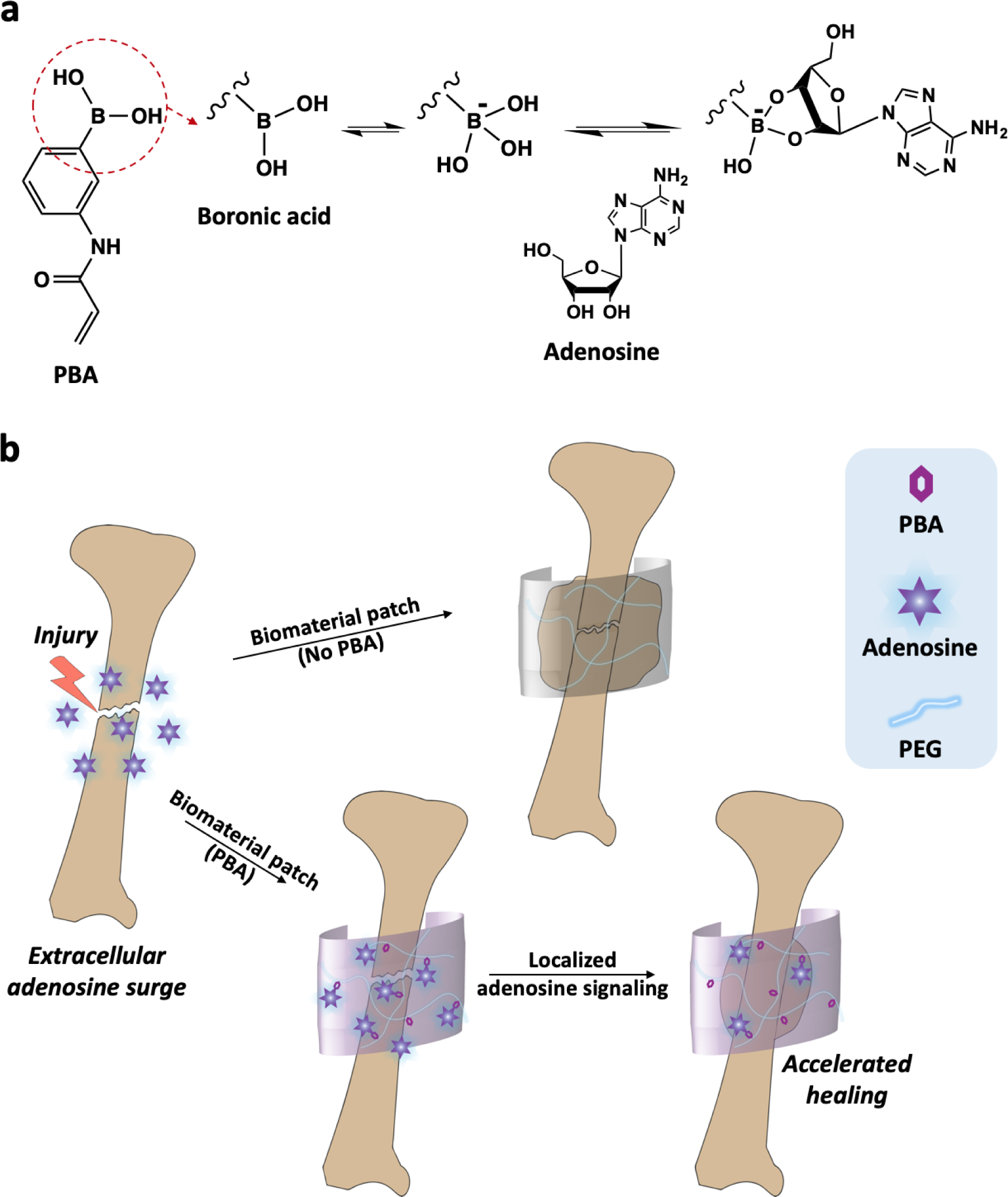
Schematics of PBA-mediated adenosine sequestration. ***a,*** 3-(acrylamido)phenylboronic acid (PBA) contains boronic acid moiety (circled in red), which forms a dynamic covalent complex of cyclic boronate ester with cis-diol-bearing adenosine at physiological pH. ***b,*** PBA-based biomaterial patch sequesters extracellular adenosine at the fracture site while leveraging the adenosine surge after injury and sustains a localized adenosine signaling to accelerate tissue repair.

## Results and Discussion

### Scaffolds functionalized with PBA groups sequester adenosine both *in vitro* and *in vivo*

To examine the PBA-assisted sequestration and release of adenosine, we have created macroporous PEG scaffolds containing varying amounts of PBA (0, 0.5 M, and 1 M as in the reaction mixture), termed as PBA_0_, PBA_0.5_, and PBA_1.0_, respectively. The macroporous PEG scaffolds were developed by using polymethyl methacrylate (PMMA) microspheres as a porogen, resulting in an interconnected macroporous architecture^38,39^ (Supplementary Fig. 1). UV/vis analysis of the residual PBA in the reaction mixture and nuclear magnetic resonance (NMR) spectra of the resulting scaffolds suggest more than 90% of the PBA molecules were reacted and incorporated into the network (Supplementary Table 1 and Supplementary Fig. 2). To determine PBA-mediated adenosine sequestration, the macroporous scaffolds with different levels of PBA were incubated in an excess adenosine solution (6 mg/mL in PBS) for 6 h and the bound adenosine was measured using UV/vis spectroscopy. As shown in **Fig. 2a**, the amount of sequestered adenosine increased as the amount of PBA within the scaffold increased. Specifically, the PBA_1.0_ scaffolds had a sequestration efficiency (the amount of PBA moieties involved in adenosine binding) of 75% with a loading capacity (weight percentage of adenosine in the scaffold) of 28%, while those of the PBA_0.5_ scaffolds were 59% and 11%, respectively (**Fig. 2b**). On the contrary, the PEG scaffolds without PBA moieties (*i.e.* PBA_0_) had no detectable adenosine content, suggesting that the loading of adenosine was primarily due to the PBA moieties. Since the PBA_1.0_ scaffolds sequestered more adenosine compared to PBA_0.5_, they were used for the rest of the studies. The release of adenosine was tested by incubating the PBA_1.0_ scaffolds in a cell culture medium depleted of nucleosides, which showed a robust release during the first 10 d followed by a plateau (**Fig. 2c,d**).

**Figure 2.**
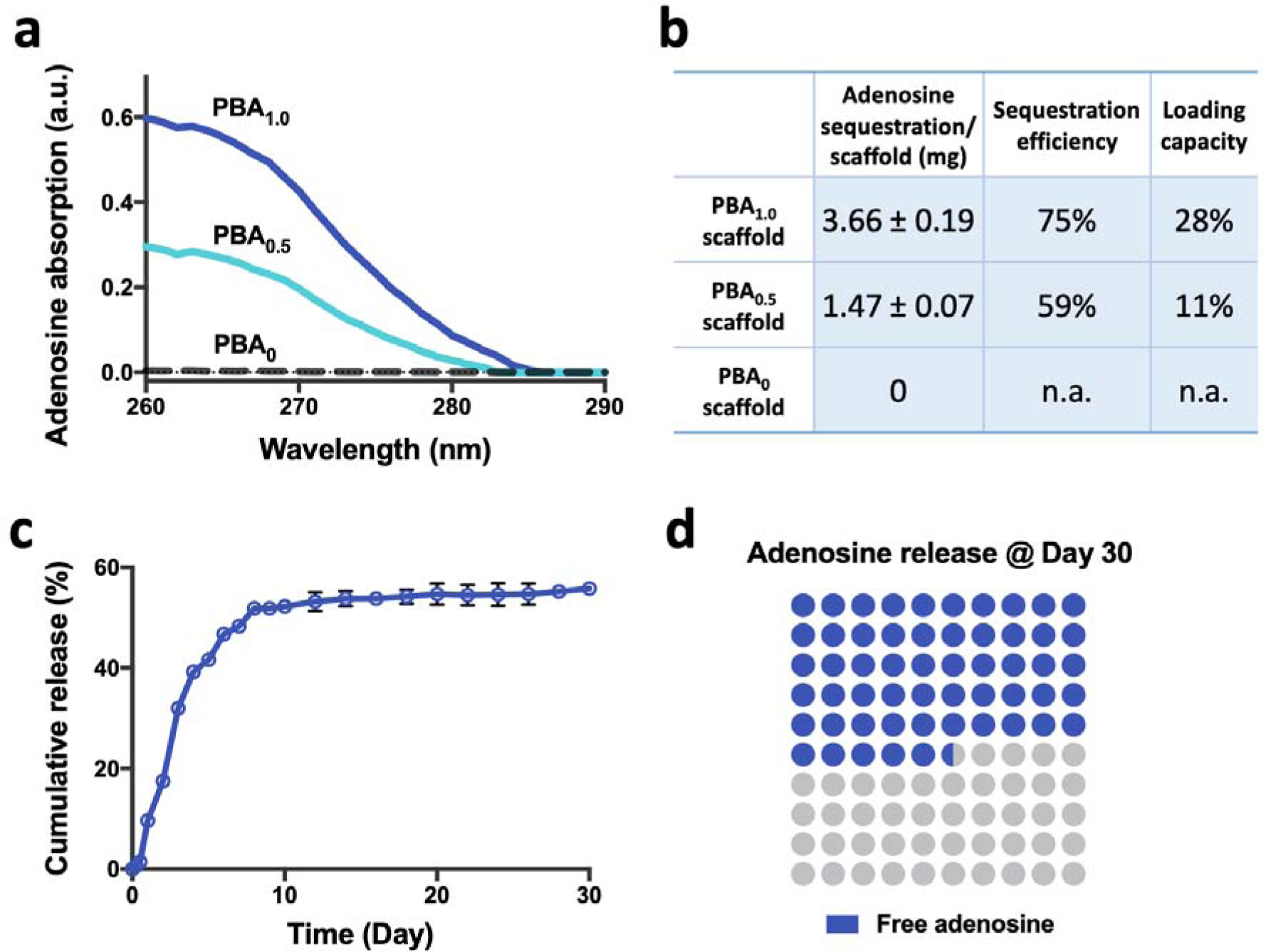
Adenosine molecules are sequestered by and released from PBA scaffolds *in vitro*. ***a,*** Representative UV/vis spectra show the absorption intensity of adenosine (in arbitrary units, a.u.), each corresponding to the amount of adenosine sequestered by a scaffold. Gray: adenosine sequestered by PBA_0_; Cyan: adenosine sequestered by PBA_0.5_; Blue: adenosine sequestered by PBA_1.0_. ***b,*** Table lists the amount of adenosine sequestered by each scaffold and the corresponding sequestration efficiency and loading capacity (n=5 scaffolds for each group). ***c,*** Cumulative release of adenosine from the PBA_1.0_ scaffolds incubated in culture medium over 30 d (n=3 PBA_1.0_ scaffolds). ***d,*** Dot map comprises blue dots representing the adenosine released from the PBA_1.0_ scaffolds by Day 30 and gray dots representing the remaining adenosine in the scaffolds. In ***b*** and ***c***, data are presented as means (± s.d.).

Having established the ability of PBA scaffolds to sequester and release adenosine *in vitro*, we next investigated their potential to sequester adenosine *in vivo*. The ability of PBA_1.0_ scaffolds to sequester adenosine *in vivo* was first assessed by using a subcutaneous model. Roughly, 600 μL of sterile saline solution containing varying amounts of adenosine (0, 0.25 or 0.5 mg/mL) was injected into an area adjacent to the scaffolds, which had been implanted subcutaneously into mice for 1 d. The scaffolds were retrieved within 1 h and analyzed for the sequestered adenosine. As anticipated, the PBA_1.0_ scaffolds retrieved from the cohort injected with 0.5 mg/mL adenosine had higher adenosine content compared to that received 0.25 mg/mL adenosine or saline alone (**Fig. 3a,b**). The PBA_1.0_ scaffolds from the cohort that received only the saline injection were also positive for adenosine, albeit a small amount, which is attributed to the endogenous adenosine present at the site of implantation. On the contrary, no adenosine was detected in the PBA_0_ scaffolds, further corroborating the necessity of PBA moieties for adenosine sequestration.

**Figure 3.**
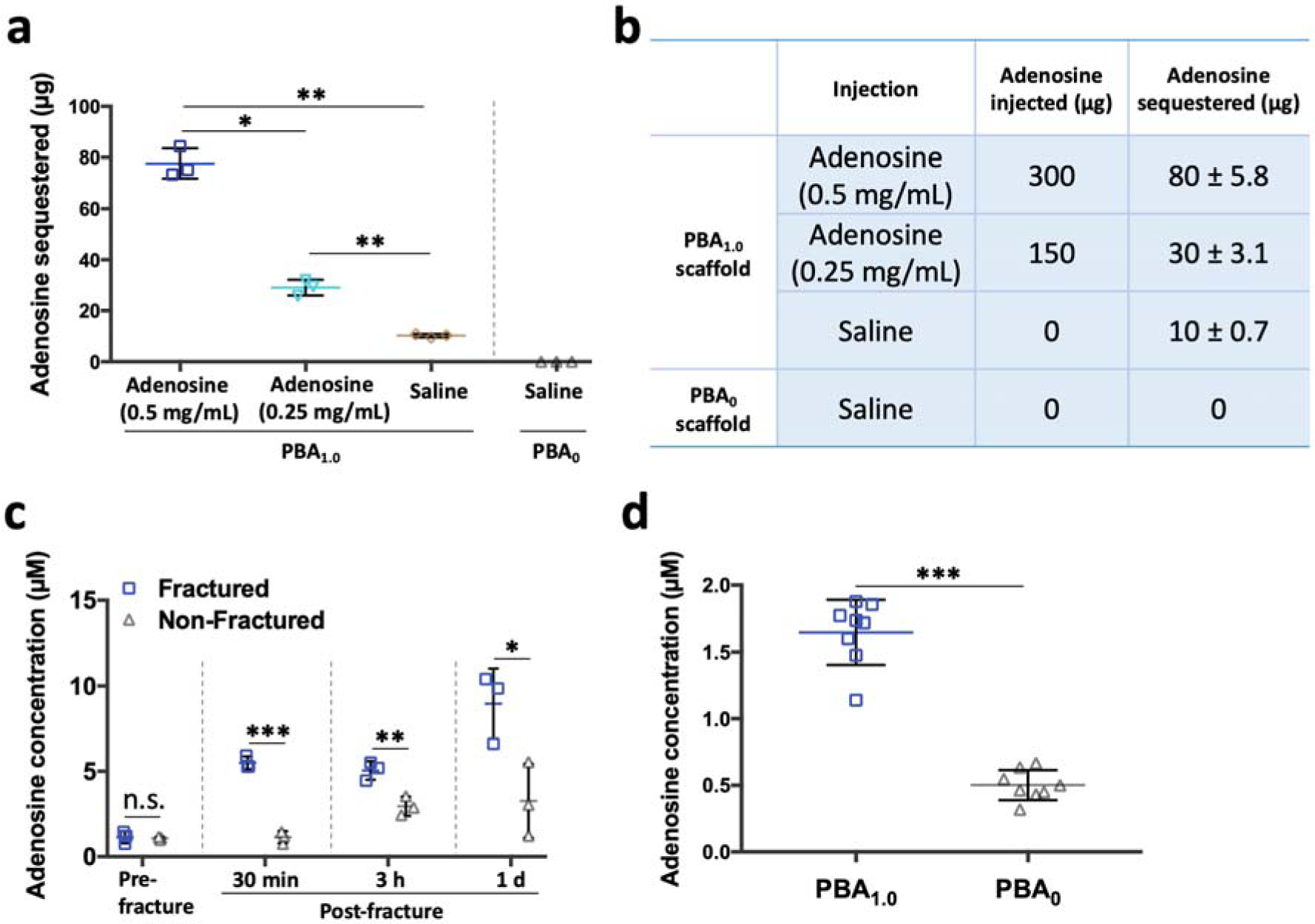
PBA scaffolds sequester adenosine *in vivo*. ***a,*** Amount of adenosine sequestered in each subcutaneously implanted scaffold following the injection of saline, 0.25 mg/mL and 0.5 mg/mL adenosine solution, respectively (n=3 scaffolds for each injection condition). ***b,*** Table lists the injection conditions, the amount of adenosine injected, and the amount of adenosine sequestered *in vivo* by each scaffold (n=3 scaffolds for each injection condition). ***c,*** Extracellular adenosine level in bone marrow before and after the unilateral fracture. Bone marrow contents from both the fractured limb and the non-fractured limb were tested (n=3 bone marrow specimens for each condition). ***d,*** Scaffolds were excised from the fracture site at 3 d post implantation, and the sequestered adenosine was quantified (n=8 scaffolds for each group). All data are presented as means (± s.d.). One-way ANOVA with Tukey’s multiple-comparisons test was used for statistical analysis in ***a***; a two-tailed *t*-test (unpaired) was used for ***c*** and ***d***. Significance is determined as \**P* < 0.05, \*\**P* < 0.01, \*\*\**P* < 0.001, and n.s. (not significant).

Although the physiological extracellular adenosine concentration in most organs is low, its level in the extracellular milieu is known to increase following trauma or injury^21,22^. Consistent with the existing knowledge, our time-dependent analyses of extracellular adenosine following unilateral tibial fracture of mouse showed a significant increase in the adenosine level at the injury site compared to that at the non-fractured contralateral site (**Fig. 3c**). A roughly 10-fold increase in extracellular adenosine was observed within 1 d following the injury. To determine the ability of PBA scaffolds to sequester extracellular adenosine by leveraging its surge following fracture, scaffolds were implanted at tibial fracture site upon injury and excised after 3 d. Compared to the PBA_0_ scaffolds retrieved from the fracture site, those as-retrieved PBA_1.0_ scaffolds contained a significantly higher amount of adenosine (**Fig. 3d**). This increased adenosine content within the implant was found to be diminished to a concentration similar to pre-fracture levels by 21 d (Supplementary Fig. 3). Together, the results suggest that biomaterials containing PBA molecules can be used to sequester and enrich extracellular adenosine locally in response to injury.

### PBA-Adenosine conjugation promotes stem cell osteogenesis both *in vitro* and *in vivo*

The osteoanabolic potential of adenosine bound to the scaffold was examined *in vitro* in a 3D culture by using human mesenchymal stem cells (hMSCs) as a cell source. Towards this, macroporous scaffolds with and without adenosine (PBA_1.0_-ADO and PBA_1.0_, respectively) were loaded with hMSCs and cultured in growth medium (GM). Cell-laden PBA_1.0_ scaffolds cultured in osteogenic-inducing medium (OM) were used as a positive control. We have previously shown that macroporous scaffolds with an interconnected macroporous structure can facilitate infiltration of the loaded cells, allowing their homogenous distribution within the scaffold^38,39^. PicoGreen DNA assay as a function of culture time showed comparable levels of DNA content in all groups (Supplementary Fig. 4). Osteogenic differentiation of hMSCs in various culture conditions was evaluated through time-resolved quantitative analyses for multiple osteogenic genes − osteocalcin (OCN), osteopontin (OPN) and osterix (OSX). As shown in **Fig. 4a**, the expressions of OCN, OPN, and OSX were consistently up-regulated throughout 21 d of culture in the PBA_1.0_-ADO scaffolds similar to the positive control. In contrast, the expressions of osteogenic markers remained low in corresponding cultures with scaffolds lacking adenosine. Consistent with these findings, quantification of calcium content exhibited significantly higher calcium deposition in the PBA_1.0_-ADO scaffolds compared to the PBA_1.0_ scaffolds with the same culture condition at the end of 21 d (**Fig. 4b**). These results suggest that the sequestered adenosine within the scaffolds promoted osteogenic differentiation of hMSCs akin to cultures involving medium supplemented with adenosine^17,35^.

**Figure 4.**
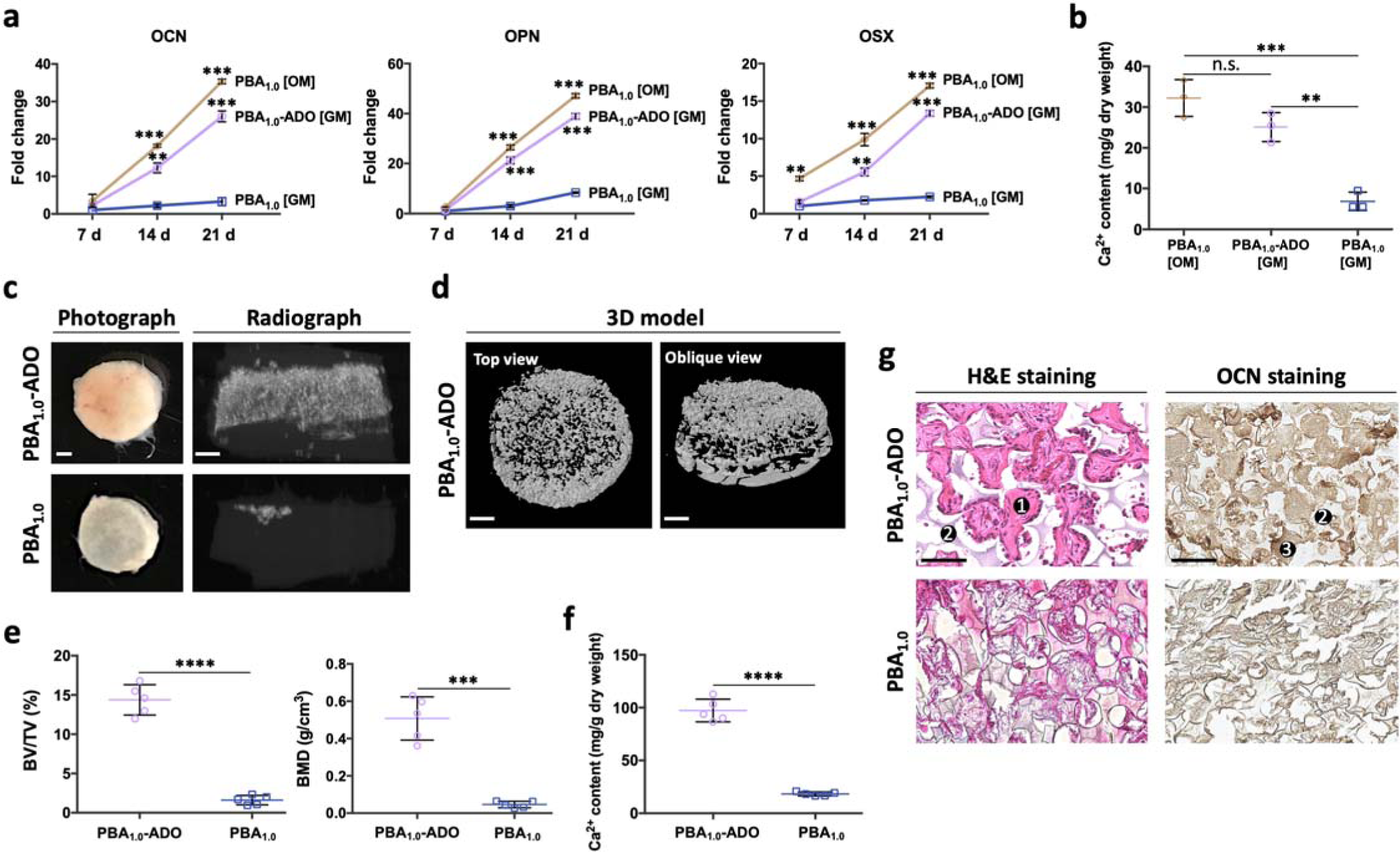
Adenosine-sequestered PBA scaffolds support osteogenic differentiation of hMSCs both *in vitro* and *in vivo*. ***a,*** PBA_1.0_ and PBA_1.0_-ADO scaffolds loaded with hMSCs were cultured *in vitro* for 21 d. Expression levels of osteogenic markers (OCN, OPN, and OSX) were quantified as a function of time and presented as fold change against 18s levels (n = 3 purified gene specimens for each group). GM: growth medium; OM: osteogenic-inducing medium. ***b,*** Calcium content in each scaffold at 21 d (n = 3 scaffolds for each group). ***c,*** Representative photos and corresponding radiographs of the cell-laden PBA_1.0_ and PBA_1.0_-ADO scaffolds excised at Day 28 post subcutaneous implantation (n = 5 scaffolds for each group). Scale bars, 1 mm. ***d,*** 3D reconstruction of a retrieved cell-laden PBA_1.0_-ADO scaffold shows the distribution of calcification, presented in both top view and oblique view. Scale bars, 1 mm. ***e,*** Bone volume ratio (BV/TV) and bone mineral density (BMD) of the retrieved scaffolds were quantified based on microcomputed tomography (n = 5 scaffolds for each group). ***f,*** Calcium content in each scaffold at Day 28 (n = 5 scaffolds for each group). ***g,*** Representative microscopic images of H&E staining and OCN immunohistochemical staining within the retrieved scaffolds (n = 5 scaffolds for each group). 1, neo-bone tissue; 2, scaffold; 3, OCN-positive tissue. Scale bars, 100 μm. All data are presented as means (± s.d.). One-way ANOVA with Tukey’s multiple-comparisons test was used for statistical analysis in ***b***; a two-tailed *t*-test (unpaired) was used for ***a*** (with reference to the group of PBA_1.0_ [GM] at 7 d)*, **e**,* and ***f***. Significance is determined as \*\**P* < 0.01, \*\*\**P* < 0.001, \*\*\*\**P* < 0.0001, and n.s. (not significant).

We next evaluated the potential of adenosine-bound scaffolds to support *in vivo* bone formation by adopting an ectopic model^40^. Both PBA_1.0_-ADO and PBA_1.0_ scaffolds loaded with hMSCs were implanted into the subcutaneous space of immunodeficient mice for 28 d. Upon retrieval, the PBA_1.0_-ADO scaffolds were found to be opaque (**Fig. 4c** and Supplementary Fig. 5). Radiographs generated from the microcomputed tomography (μCT) scans showed a strong optical signal from the PBA_1.0_-ADO scaffolds, which is consistent with the gross appearance, suggesting *in vivo* calcification and the presence of hard tissue formation (**Fig. 4c**). The 3D rendering of the excised PBA_1.0_-ADO scaffolds showed an even distribution of mineral deposition within the scaffolds, as indicated by both top and oblique views (**Fig. 4d**). Conversely, the excised PBA_1.0_ scaffolds did not display this opaque appearance nor apparent calcification. Based on the quantification of μCT results, the PBA_1.0_-ADO scaffolds had a bone volume ratio (BV/TV) of 14.4% and a bone mineral density (BMD) of 0.51 g/cm^3^, compared to 1.6% and 0.05 g/cm^3^ found within the PBA_1.0_ group (**Fig. 4e**). Measurement of calcium content within the scaffolds, 97.3 ± 4.8 mg/g dry weight in the PBA_1.0_-ADO and 18.2 ± 0.8 mg/g dry weight in the PBA_1.0_ scaffolds (**Fig. 4f**), further confirmed higher *in vivo* calcification of the cell-laden PBA_1.0_-ADO scaffolds.

Bone tissue formation was further evaluated by histological characterization. Hematoxylin and eosin (H&E) staining of the excised implants showed dense extracellular matrix (ECM), resembling that of the bone tissue, in the cell-laden PBA_1.0_-ADO scaffolds, whereas the corresponding PBA_1.0_ group had minimal bone tissue formation (**Fig. 4g**). Furthermore, positive staining of OCN, an ECM protein secreted by osteoblasts, was seen throughout the PBA_1.0_-ADO scaffolds (**Fig. 4g**). Together, the findings suggest that the adenosine-loaded scaffolds supported osteogenic differentiation of the transplanted hMSCs and promoted ectopic bone formation, which further corroborates the osteoblastogenic function of adenosine.

### PBA-mediated adenosine sequestration promotes bone fracture healing

We employed a tibial fracture model to investigate the role of biomaterial-assisted sequestration of adenosine in bone repair, which is a comprehensive process involving cartilaginous callus formation at the injury site, endochondral ossification within the callus, and callus/bone remodeling^41^. Stabilized fractures were induced unilaterally at tibial midshafts in mice^42^, and biomaterials with uniform dimensions were used to cover the fracture sites. In addition to PBA_0_ and PBA_1.0_ patches, we also used patches pre-loaded with exogenous adenosine (PBA_1.0_-ADO). We monitored the fracture healing as a function of time using radiographic and histomorphometric analyses.

**Figure 5a** shows 3D reconstructed images (intact and cut views) and their corresponding radiographs of the injured tibiae during fracture healing. The evolution of callus as bone repair progresses is represented in **Figure 5b**. At 7 d, the fractures were still evident in all groups owing to the minimal mineralization of the calluses. As time progressed, the calluses calcified, and the extent of mineralization was found to correlate with the type of intervention, where both the PBA_1.0_-ADO and the PBA_1.0_ groups exhibited a better bridging of the fracture. By 21 d, growing calluses eventually bridged the fracture gaps in all groups. Interestingly, radiographic images at 21 d showed cortical bridging only in groups treated with PBA_1.0_-ADO and the PBA_1.0_ (**Figure 5a,b**), suggesting faster healing compared to those treated with PBA ^41^. Concomitant with these observations, analysis of the fracture sites at 21 d from the axial view (**Figure 5c**) revealed better remodeled patterns and more organized lamellar bone formation in cohorts that received either PBA_1.0_-ADO or PBA_1.0_ patches. These findings were further confirmed by the quantification of the μCT scans at 14 d (**Figure 5d**) and 21 d (**Figure 5e**). By 14 d, bone volume was higher in both the PBA_1.0_-ADO and the PBA_1.0_ groups, albeit with no statistical significance. The differences in bone formation were apparent by 21 d, where the fractures treated with PBA_1.0_-ADO and PBA_1.0_ exhibited significantly higher bone volume ratio within the calluses compared to those treated with PBA_0_. Together, the results demonstrate the prevalent role of localized adenosine signaling in promoting callus maturation and fracture healing. When the biomaterial patch was dosed once with exogenous adenosine, as in the case of the PBA_1.0_-ADO group, the healing was further improved, mostly due to the higher amount of adenosine available.

**Figure 5.**
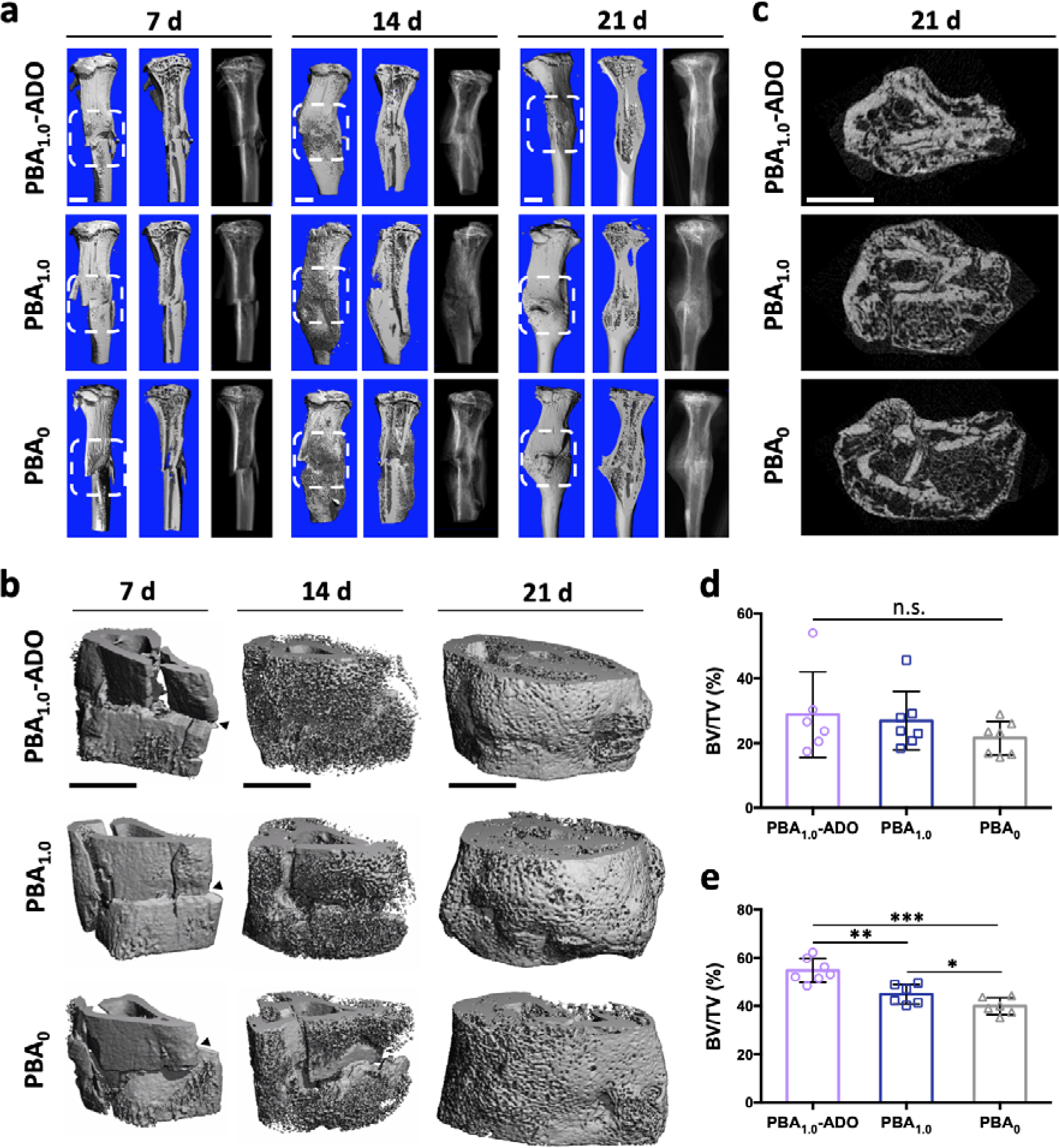
PBA-containing biomaterial patches promote callus maturation during fracture healing. ***a,*** 3D reconstructions (intact and cut views) and corresponding radiographs of the fractured tibiae treated with various biomaterial patches. Tissues were harvested at 7 d, 14 d, or 21 d following injury (n = 7 mice for each treatment). White boxes cover the callus regions. Scale bars, 1 mm. ***b,*** Magnified 3D reconstructions of the callus regions show the evolution of calluses during fracture healing with black arrowheads indicating fracture sites. Scale bars, 1 mm. ***c,*** Representative axial views of fracture sites show remodeling outcome within calluses by 21 d. Scale bar, 1 mm. ***d,*** Bone volume ratio (BV/TV) of calluses at 14 d was quantified based on microcomputed tomography (n = 6 mice for the cohort treated with PBA_1.0_-ADO, n = 7 mice for the cohort treated with PBA_1.0_ or PBA_0_). ***e,*** Bone volume ratio (BV/TV) of calluses at 21 d was quantified (n = 7 mice for the cohort treated with PBA_1.0_-ADO, n = 6 mice for the cohort treated with PBA_1.0_ or PBA_0_). In ***d*** and ***e***, data are presented as means (± s.d.). One-way ANOVA with Tukey’s multiple-comparisons test was used for statistical analysis in ***d*** and ***e***. Significance is determined as \**P* < 0.05, \*\**P* < 0.01, \*\*\**P* < 0.001, and n.s. (not significant).

Given the importance of the evolution of cartilaginous tissue, vascularization, and osteoclast-driven bone resorption in fracture healing^41^, we also examined the effect of biomaterial-mediated adenosine signaling on cartilaginous tissue, blood vessel formation, and osteoclast activity during healing. We observed intense cartilage formation (stained red) within the calluses of both the PBA_1.0_-ADO and the PBA_1.0_ groups at 7 d (**Fig. 6a**), followed by cartilage resorption over time suggesting endochondral ossification. In contrast, the animals treated with the PBA_0_ showed delayed cartilaginous tissue formation and remodeling. Cartilaginous tissues still remained in the calluses of these animals at 21 d (**Fig. 6a**). Concurrent with these findings, an intervention-specific change in vascularization of the calluses was also observed (**Fig. 6b-e**). Specifically, more endomucin (EMCN)-positive blood vessels were detected in both the PBA_1.0_-ADO and the PBA_1.0_ groups compared to the PBA_0_ cohort at 7 d (**Fig. 6c**) and 14 d (**Fig. 6d**). The improved callus vascularization in the presence of PBA-mediated adenosine signaling may be directly linked to the established role of adenosine in promoting angiogenesis^43,44^. The improved osteoblastogenesis observed in these groups could also contribute to the increased angiogenesis^45,46^. While there were differences in angiogenesis among the different groups at early time points, no intervention-dependent differences in blood vessel content were observed at 21 d (**Fig. 6e**). Analyses of osteoclast activity via TRAP staining (Supplementary Fig. 6a) showed increased TRAP-positive area with time in all the groups. Particularly, higher percentage of TRAP-positive area in the PBA_1.0_-ADO cohort at 21 d (Supplementary Fig. 6b), indicating greater bone remodeling, which could be associated with high levels of bone formation.

**Figure 6.**
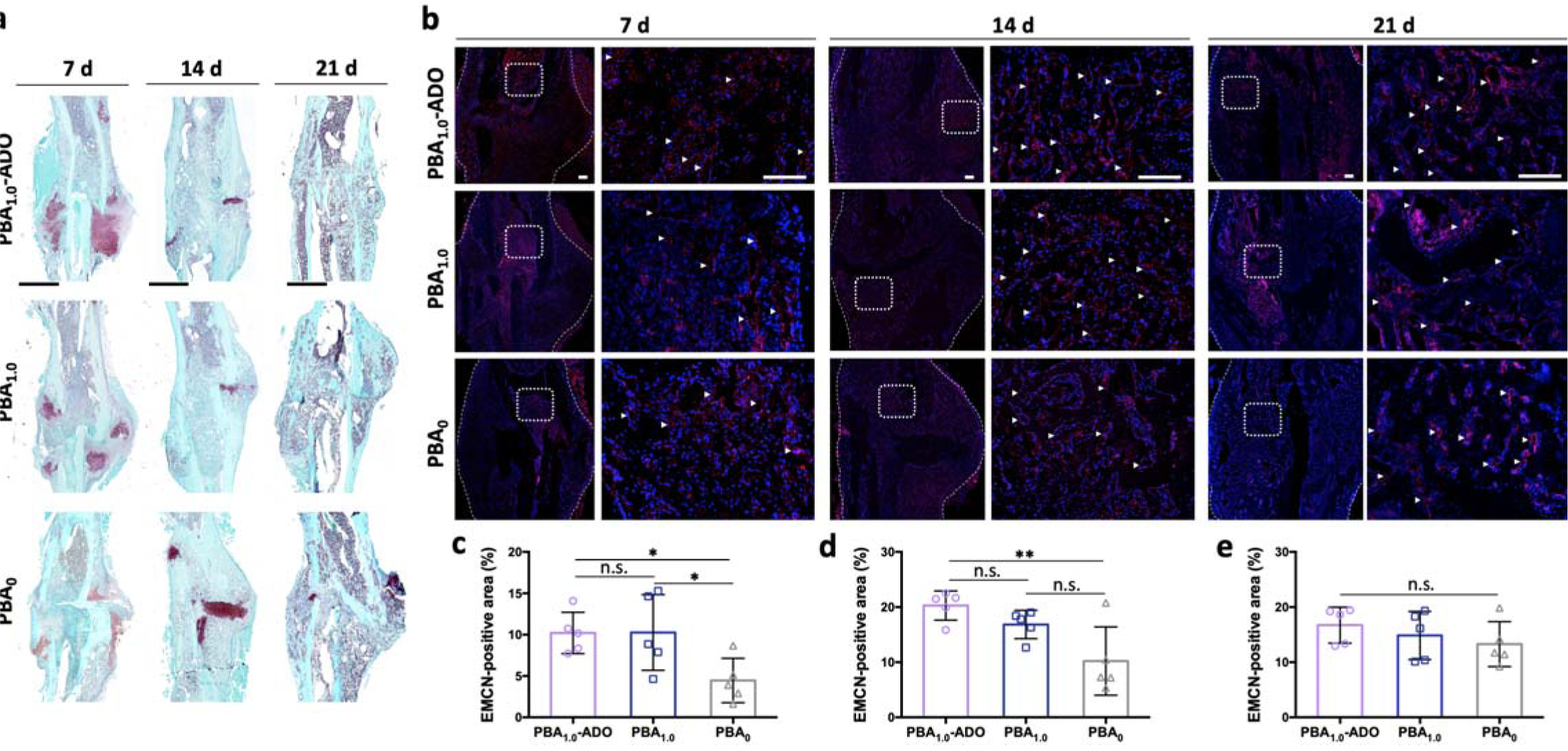
PBA-containing biomaterial patches facilitate endochondral ossification and vascularization in fracture calluses. ***a,*** Representative safranin-O staining images of fractured tibiae treated with various biomaterial patches show the evolution of cartilaginous tissues (red) in calluses (n = 7 mice for each treatment). Scale bars, 1 mm. ***b,*** Representative immunofluorescence images of endomucin (EMCN, red) show the vascularization in calluses (n = 7 mice for each treatment). Nuclei were stained with DAPI (blue). White dashed lines delineate the callus boundary. Magnified images of the region covered in white boxes are displayed with white arrowheads indicating EMCN-positive vessels. Scale bars, 200 μm. ***c,d,e,*** Quantification of the relative EMCN-positive vessel area to the callus area at 7 d (***c***), 14 d (***d***), and 21 d (***e***) based on immunofluorescence images (n = 5 mice from each treatment; each data point is averaged from 5 images for each mouse). In ***c***-***e***, data are presented as means (± s.d.). One-way ANOVA with Tukey’s multiple-comparisons test was used for statistical analysis in ***c***-***e***. Significance is determined as \**P* < 0.05, \*\**P* < 0.01, and n.s. (not significant).

To summarize, the cohorts treated with PBA_1.0_ or PBA_1.0_-ADO showed a better healing outcome compared to those treated with PBA_0_, as evidenced by the extent of callus maturation, endochondral calcification, and angiogenesis, suggesting that biomaterial-mediated localization of adenosine signaling promotes bone healing. Furthermore, the increased adenosine concentration from the biomaterial-mediated sequestration recedes to the physiological level with healing (Supplementary Fig. 3), which underscores the translational potential of the described strategy. While PBA-mediated sequestration of endogenous adenosine alone promoted fracture healing, the augmentation of adenosine level with a one-time supplement of exogenous adenosine (*i.e.* PBA_1.0_-ADO) further improved callus vascularization and healing outcome. Note that while the PBA_1.0_ sequestered only a small fraction of the adenosine being released by cells (**Fig. 3d**), further improvement of biomaterial design, such as increasing PBA content or changing the architecture to increase surface-to-volume ratio, could be used to enhance the sequestration efficiency. Such an approach will imbibe more adenosine from the milieu following injury and sustain its local concentration and could eliminate the need for exogenous adenosine entirely. Nonetheless, the results presented in this study showed the potential of using a PBA-containing biomaterial to boost the adenosine concentration at the fracture site and leverage the natural repair mechanism involving adenosine signaling to promote fracture healing.

### Conclusion and Outlook

This proof-of-concept study demonstrates that sequestration of adenosine, a native small molecule, by biomaterials at the fracture site can be used to promote bone fracture healing. The sequestration and release of adenosine was achieved by harnessing the ability of boronate molecules to form dynamic covalent bonds with cis-diol molecules such as adenosine. This biomaterial approach sustained an elevated concentration of adenosine locally by leveraging the surge of extracellular adenosine following injury and created a pro-regenerative milieu through localized adenosine signaling, resulting in improved bone repair. Besides sequestering endogenous adenosine, the biomaterial can also be used to deliver exogenous adenosine to the injury site, especially in pathological situations encountering diminished extracellular adenosine. By enabling prolonged adenosine signaling, this biomaterial approach circumvents potential off-target effects associated with the systemic administration of adenosine, which is a major hurdle in harnessing adenosine signaling as a potential therapeutic.

Moving forward, the biomaterial-assisted sequestration of extracellular adenosine can be used to create an in-situ stockpile of the small molecule, which can be conveniently replenished non-invasively through injections. For example, local modulation of the adenosine signaling may be used to prevent repeated fractures, which are commonly observed in the aged population, as well as in patients suffering with osteoporosis and other bone-degenerating diseases. In the future, age- and sex-dependent fracture healing in response to extracellular adenosine can also be examined to determine how the adenosine metabolism may differ and lead to different repair outcomes. The biomaterial can be adapted accordingly to mirror the innate repair mechanism by modulating the extent of adenosine sequestration.

### Experimental Section

#### Materials

Polyethylene glycol diacrylate (PEGDA) and N-acryloyl-6-aminocaproic acid (A6ACA) were synthesized as previously described^47,48^. Briefly, 10 wt% of PEG (Mw∼3.4 kDa, Sigma-Aldrich, Cat# P4338) solution was made by dissolving it in anhydrous dichloromethane (DCM) at room temperature with stirring in an argon environment. To this, 1.5 molar equivalents of triethylamine (Sigma-Aldrich, Cat# 471283) and acryloyl chloride (Sigma-Aldrich, Cat# A24109) were added dropwise on ice. The reaction was continued overnight at room temperature followed by purification using Celite diatomaceous earth (Sigma-Aldrich, Cat# 1026931000). The product was concentrated using a rotary evaporator and precipitated in ∼10-fold chilled diethyl ether. The resultant PEGDA was dried under vacuum overnight and purified by using a Sephadex fine G-25 column (GE Healthcare Life Sciences), followed by lyophilization. To synthesize A6ACA, 1 M of 6-aminocaproic acid (Sigma-Aldrich, Cat# A7824) was prepared by dissolving it in 1N sodium hydroxide, followed by reacting with 1.5 molar equivalent of acryloyl chloride added dropwise on ice. The reaction mixture was maintained at pH 8 for an hour and then gradually decreased to 3 by titrating with 5N hydrochloric acid. The product was extracted using ethyl acetate, dried over anhydrous sodium sulfate, and precipitated in chilled hexane. The resultant A6ACA was collected and dried overnight under vacuum. The successful completion of the reactions was confirmed by NMR as described previously^48^.

#### Macroporous scaffold fabrication

PEG macroporous scaffolds containing the PBA moieties were fabricated using a poly(methyl methacrylate) (PMMA) leaching method^38^. The polymer precursor solution was prepared by mixing PEGDA (10% w/v), 3-(acrylamido)phenylboronic acid (PBA, 1 M or 0.5 M; Sigma-Aldrich, Cat# 771465), A6ACA (0.5 M) and Irgacure 2959 (0.5% w/v; Sigma-Aldrich, Cat# 410896) in 80% ethanol. 20 μL of the precursor solution was added into a cylindrical polypropylene mold (5 mm in diameter) packed with 20 mg of PMMA microspheres (150-180 μm, Cospheric), followed by UV irradiation (365 nm) for 10 min. The resultant structures were soaked in acetone for 3 d to remove the PMMA beads with daily change of solvent followed by washing with deionized water. The macroporous scaffolds were trimmed to a uniform size of 6 mm in diameter and 2 mm in height. PEG macroporous scaffolds without PBA were also prepared in the same way. For sterilization, the scaffolds were soaked in 70% ethanol for 6 h, followed by washing in sterile PBS extensively for 3 d. The sterilized scaffolds were used for cell loading and *in vivo* experiments.

#### Scanning Electron Microscopy (SEM)

Scaffolds were cut into thin slices and freeze-dried. To examine their porous structure, sliced scaffolds were sputter-coated with Au for 100 s (Denton Desk IV) and imaged by using a Philips XL30 ESEM under high vacuum mode.

#### Nuclear magnetic resonance (NMR) spectroscopy

To examine the extent of PBA incorporation, freshly prepared PBA scaffolds were thoroughly washed with deionized water to remove the unreacted PBA and freeze-dried. The samples were then minced and fully solvated in heavy water (D_2_O) as described earlier^48^. PBA_0_ scaffolds were also prepared the same way. ^1^H NMR spectra were recorded for all the samples using a 400 MHz Varian Inova spectrometer.

#### Adenosine loading and release

Macroporous scaffolds were soaked in PBS supplemented with 6 mg/mL adenosine (Sigma-Aldrich, Cat# A4036) for 6 h and washed thoroughly to remove unbound adenosine. To measure the amount of sequestered adenosine, the scaffolds were soaked in acetate buffer (0.1 M, pH 4.5) for 2 h to completely release adenosine into the buffer, which was subsequently analyzed by using a UV/vis spectrophotometer (Beckman Coulter) at a wavelength of 260 nm. The concentration of the released adenosine was determined from a standard curve generated using adenosine solutions with known concentrations, ranging from 0.5 mM to 5 mM. To characterize the release profile of adenosine, the scaffolds were incubated in PBS with or without glucose (50 mM), or incubated in αMEM (Gibco, Cat# 12561056) containing 10% (v/v) fetal bovine serum at 37 °C. The concentration of adenosine in the buffer or medium was monitored as a function of time through UV/vis spectrophotometry.

#### Cell culture and loading

Primary hMSCs were maintained and expanded in growth medium (GM) containing high-glucose DMEM (Gibco, Cat# 11995065), 10% (v/v) fetal bovine serum (HyClone, Cat# SH3007103HI), 4 mM L-glutamine (Gibco, Cat# 35050061), and 50 U/mL Penicillin/Streptomycin (Gibco, Cat# 15140122). Cells were passaged at 70-80% confluency and used at Passage 5. Prior to cell loading, the sterilized scaffolds with and without adenosine were equilibrated in growth medium at 37°C for 1 d. Cell loading was performed according to a published method^40^. Briefly, 20 µL of cell suspension containing 1 million hMSCs was loaded onto partially dehydrated scaffolds. The cell-laden scaffolds were kept in GM at 37°C for 1 d to allow for cell infiltration. For *in vivo* study, these cell-laden scaffolds were then implanted subcutaneously in mice for 28 d. For *in vitro* study, the cell-laden scaffolds were cultured in GM supplemented with 3 mM phosphate at 37°C and 5% CO_2_ with medium change every other day. As a positive control for *in vitro* osteogenesis, a group of cell-laden scaffolds without adenosine were cultured in osteogenic-inducing medium (OM) made of GM supplemented with 10 mM β-glycerophosphate (Sigma, Cat# G9422), 50 µM ascorbic acid (Sigma, Cat# A4403), and 100 nM dexamethasone (Sigma, Cat# D2915)^49^.

#### Cell viability assay

PBA_1.0_ and PBA_1.0_-ADO scaffolds loaded with hMSCs were cultured *in vitro* in different culture media (GM: growth medium, OM: osteogenic-inducing medium) for 21 d. The DNA content in the scaffolds as a function of time was quantified following a fluorescence method^50^. The scaffolds were thoroughly washed in PBS and homogenized in 0.2% Triton X-100. The lysate was then diluted as needed and combined with a same volume of 1X PicoGreen dsDNA reagent in 1X TE buffer provided in the PicoGreen assay kit (Invitrogen, Cat# P11496). After 5 min, the fluorescence intensity of the mixture at 480 nm/520 nm (excitation/emission) was measured. The DNA content was determined according to a standard curve of lambda DNA.

#### RNA isolation and quantitative real-time polymerase chain reaction (qRT-PCR)

Osteogenic differentiation of the hMSCs as a function of time was evaluated by using qRT-PCR. The cell-laden scaffolds were lysed and homogenized in TRIzol Reagent (Invitrogen, Cat# 15596018). Total RNA was extracted with chloroform and precipitated in isopropanol. 1 μg of each RNA sample was reverse transcribed with the iScript Reverse Transcription Supermix (Bio-Rad, Cat# 1708841) following the manufacturer’s instructions. The obtained cDNA was mixed with the forward and reverse primers of the target gene along with the iTaq SYBR Green Supermix (Bio-Rad, Cat# 1725124) for qRT-PCR according to the manufacturer’s protocol. The qRT-PCR was conducted in a Bio-Rad Thermal Cycler (CFX96) following the steps of an initial denaturation at 95°C for 30 s for 1 cycle, amplifications at 95°C for 5 s and 60 °C for 30 s for 40 cycles, and finally 95°C for 10 min. The primer sequences are: osteocalcin (OCN; forward: TGAGAGCCCTCACACTCCTC; reverse: ACCTTTGCTGGACTCTGCAC), osteopontin (OPN; forward: AATTGCAGTGATTTGCTTTTGC; reverse: CAGAACTTCCAGAATCAGCCTGTT), osterix (OSX; forward: CATCTGCCTGGCTCCTTG; reverse: CAGGGGACTGGAGCCATA), and 18s (forward: CCCTGTAATTGGAATGAGTCCACTT; reverse: ACGCTATTGGAGCTGGAATTAC). The expression level of each target gene was calculated as ΔCt relative to the corresponding housekeeping gene (18s), converted to 2^(-ΔΔCt) by normalizing to the group of PBA scaffolds cultured in growth medium for 7 days, and presented as fold change.

#### Calcium assay

To quantify the calcium deposition, cell-laden scaffolds were washed in deionized water, freeze-dried, lyophilized, and homogenized in 0.5 N HCl. The resultant homogenate was added into a Calcium Assay solution (Pointe Scientific, Cat# C7503) and the absorbance at 570 nm was recorded using a Multimode Detector^35^. The amount of free calcium ions (Ca^2+^) in the mixture was determined from a standard curve of calcium chloride solutions with known concentration and normalized to the dry weight of the corresponding scaffold.

#### Subcutaneous implantation and tibial fracture

All animal studies were conducted with the approval of the Institutional Animal Care and Use Committee (IACUC) at Duke University and complied with NIH guidelines for laboratory animal care. Female immunodeficient NOD.CB17-Prkdcscid/J mice (4-month-old, Jackson Lab) were used for subcutaneous implantation of hMSC-laden PBA scaffolds with and without adenosine. Female C57BL/6J mice (4-month-old, Jackson Lab) were used for all the experiments involving acellular scaffolds and patches. The mice were anesthetized with 2% isoflurane and administered with buprenorphine (1 mg/kg, sustained release, ZooPharm) through subcutaneous injection prior to surgical procedure. For subcutaneous implantation^40^, a roughly 1 cm-long incision was made on the back of each anesthetized mouse, and each cell-laden scaffold was implanted to the right side of the subcutaneous pouch. For tibial fracture^42^, the right tibia was first stabilized by inserting a 0.7 mm pin from the tibial plateau through the medullary cavity after removal of the skin proximal to the knee, a fracture was then created at the tibial midshaft with blunt scissors, and an 8 mm by 3 mm biomaterial patch was wrapped around the fracture site and held tight underneath the muscle. Upon completion, two drops of bupivacaine (0.5%, Hospira) were applied topically along the incision line, followed by closure with wound clips.

#### Extracellular adenosine level in bone marrow

Bone marrow specimens from both the fractured limbs and the contralateral non-fractured limbs of the mice were collected at 30 min, 3 h, and 1 d following tibial fractures, respectively. Plasma was isolated from the bone marrow flush by centrifugation at 2,000 g, 4°C for 20 min, and was subsequently diluted with an adenosine-protecting solution containing 0.2 mM dipyridamole, 5 µM erythro-9(2-hydroxy-3-nonyl)-adenine, 60 µM alpha, beta-methylene-adenosine 5’-diphosphate, and 4.2 mM ethylenediaminetetraacetic acid (EDTA). Extracellular adenosine content in the diluted plasma was quantified by using an Adenosine Assay Kit (Fluorometric; Abcam, Cat# ab211094) following the manufacturer’s instructions. Briefly, each sample was mixed with a series of reagents including Adenosine Detector, Adenosine Convertor, Adenosine Developer and Adenosine Probe in an Adenosine Assay Buffer, and the mixture was incubated in dark for 15 min. Fluorescence intensity of the mixture was measured at 535 nm (excitation)/590 nm (emission) using a Multimode Detector, and the adenosine concentration was determined based on known adenosine standards.

#### In vivo adenosine sequestration

Freshly prepared PBA_1.0_ and PBA_0_ scaffolds with identical dimensions were separately implanted into the subcutaneous pouches or the tibial fracture site of mice. For adenosine sequestration in the subcutaneous space, each mouse received a subcutaneous injection of 600 μL sterile saline or adenosine solution (0.25 mg/mL, 0.5 mg/mL) in the vicinity of the scaffolds at 1d after the subcutaneous implantation, and the scaffolds were excised 1 h later. For adenosine sequestration at the fracture sites, the scaffolds were excised at 3 d and 21 d after the implantation, respectively. The as-retrieved scaffolds were rinsed in PBS, minced, and soaked in acetate buffer (0.1 M, pH 4.5) for 2 h. The supernatant was subsequently collected, neutralized, and used for adenosine measurement.

#### Microcomputed tomography

Scaffolds retrieved from the subcutaneous implantation and fractured tibiae of mice were collected, processed in 4% paraformaldehyde at 4°C for 3 d, and rinsed with PBS. The fixed samples and a phantom were loaded into a µCT scanner (vivaCT 80, Scanco Medical) and scanned at 55 keV with a pixel resolution of 10.4 µm. Reconstruction of the scanned images was performed using µCT Evaluation Program V6.6 (Scanco Medical), followed by generation of radiographs and 3D images using µCT Ray V4.0 (Scanco Medical). Bone mass was evaluated based on the reconstructed images and presented as a percentage of bone volume over total volume (%BV/TV). Bone mineral density (BMD) was determined by using the phantom with known hydroxyapatite content as a reference. Calluses of fractured tibiae were analyzed for %BV/TV based on 200 contiguous slices within 1 mm proximal and 1 mm distal of the fracture center according to a published study^42^.

#### Histological analyses

Fixed samples were decalcified in 14% EDTA (pH 8.0) at 4°C for 5 d, rinsed in PBS, dehydrated and embedded in paraffin, and cut into 5 μm-thick sections by using a Leica rotary microtome. Prior to staining, each section was deparaffinized in CitriSolv (Decon Labs, Cat# 1601) and rehydrated through graded alcohols and deionized water. For H&E staining, rehydrated sections were immersed in hematoxylin solution (Ricca Chemical, Cat# 3536-16) for 4 min and then switched in eosin-Y solution (Ricca Chemical, Cat# 2845-16) for 1 min. For osteocalcin (OCN) immunohistochemical analysis, rehydrated sections were immersed in a blocking buffer made of PBS, 3% bovine serum albumin and 0.1% Tween-20 for 1 h, and incubated with OCN primary antibody (1:100 in blocking buffer, rabbit polyclonal; Abcam, Cat# ab93876) overnight at 4°C. After rinsing thoroughly with PBS, the sections were treated with horseradish peroxidase (HRP)-conjugated secondary antibody (1:100 in blocking buffer, HRP-donkey anti-rabbit; Jackson ImmunoResearch, Cat# 711-035-152) at room temperature for 1 h, followed by incubating in a developing solution containing 3-3’ diaminobenzidine (DAB) peroxidase substrate (Vector Laboratories, Cat# SK-4100) for 5 min to produce a brown reaction product. For Safranin-O staining of the fracture calluses, rehydrated tibia sections were immersed in 1% Safranin-O (Sigma, Cat# S8884) at room temperature for 1 h and counter-stained with 0.02% Fast Green (Sigma, Cat# F7258) and hematoxylin solution for 1 min. For tartrate-resistant acid phosphatase (TRAP) staining, rehydrated tibia sections were immersed in a 0.2 M sodium acetate buffer (pH 5.0) containing 50 mM tartaric acid (Sigma, Cat# 228729), 0.5 mg/mL naphthol AS-MX phosphate (Sigma, Cat# N5000), and 1.1 mg/mL fast red TR (Sigma, Cat# F6760) for 1 h at 37°C. After rinsing in deionized water, the sections were counterstained in Mayer’s hematoxylin solution (Sigma, Cat# MHS16) for 1 min. All the stained sections were subsequently dehydrated, covered with a mounting medium (Fisher Scientific, Cat# SP15-100), and imaged using a Keyence (BZ-X710) microscopy system.

#### Immunofluorescence imaging and vessel quantification

Rehydrated tibia sections were steam-treated in a citrate buffer (pH 6.0; Abcam, Cat# ab64236) for antigen retrieval and further immersed in blocking buffer for 1 h at room temperature. The sections were then incubated with endomucin primary antibody (1:100 in blocking buffer, rat polyclonal; Abcam, Cat# ab106100) overnight at 4°C, followed by addition of secondary antibody (1:200 in blocking buffer, Alexa Fluor 647-rabbit anti-rat; Abcam, Cat# ab169349) at room temperature for 1 h. All the sections were subsequently rinsed in PBS and mounted with an antifade medium containing DAPI (Invitrogen, Cat# P36971). Images were acquired using a Zeiss (Axio Imager Z2) microscopy system under the same exposure time for all groups. Quantification of blood vessel formation in calluses was conducted by using ImageJ (v1.52g) and represented as the percentage of EMCN-positive vessel area (red) to the callus area (blue) based on the immunofluorescence images. Ten images from each mouse tibia were used, and five mice from each group were included for the analysis.

#### Statistical analysis

The means with standard deviations (n ≥ 3) are presented in the results. All the data were subjected to either two-tailed Student’s t-test or one-way analysis of variance (ANOVA) with post hoc Tukey-Kramer test for multiple comparisons using GraphPad Prism 7. Any *P*-value of less than 0.05 was indicated with asterisk and considered statistically significant.

## Associated content

Supplementary Information

## Author information

Corresponding Author: *

Notes: The authors declare no competing financial interest.

## Acknowledgments

We thank Dr. Jiaul Hoque for assisting the NMR characterization. We are grateful to Dr. Matthew Hilton, Dr. Rong Huang, and Anthony Mirando for their help with histomorphometric analysis. This work was partially supported by the National Institute of Arthritis and Musculoskeletal and Skin Diseases of the National Institutes of Health under Award Number NIH R01 AR063184 and AR071552. The content is solely the responsibility of the authors and does not necessarily represent the official views of the National Institutes of Health. The hMSCs were provided by Institute for Regenerative Medicine (Texas A&M University) through NIH Grant P40RR017447 from the National Center for Research Resources.

## Author contributions

Y.Z. and S.V. designed the study and wrote the manuscript. Y.Z. executed the study and organized the results. G.S.B and Y.V.S. conducted the animal surgeries. Y.V.S. contributed to overseeing the experiments and interpreting the data.

