## Supplementary material for "*In vivo* sequestration of innate small molecules to promote bone healing": Supplmentary Material

**Scaffold preparation and adenosine sequestration efficiency**

Due to the low solubility and low reactivity of 3-(acrylamido)phenylboronic acid (PBA) in aqueous solution^1^, we used an ethanol/water mixture as solvent and supplemented N-acryloyl-6-aminocaproic acid in the precursor solution prior to photopolymerization. To evaluate the amount of PBA being incorporated into the copolymer, the remaining PBA after photopolymerization was collected and measured at 264 nm using UV/vis spectroscopy. The spectroscopic analyses showed that 91.5% of the PBA in the precursor solution were incorporated into the PBA_1.0_ scaffold, and the PBA incorporation efficiency for the PBA_0.5_ scaffold was 93.3% (**Table 1**). Furthermore, the addition of PBA did not affect the formation of interconnected porous structure based on the examination of SEM images (**Figure 1**).

**Table 1.**

|  | PBA incorporation efficiency | Theoretical ADO sequestration (mg) | Actual ADO sequestration (mg) | ADO sequestration efficiency | ADO loading capacity |
| --- | --- | --- | --- | --- | --- |
| PBA_1.0_ scaffold | 91.5% | 5.34 | 3.66 | 75% | 28% |
| PBA_0.5_ scaffold | 93.3% | 2.67 | 1.47 | 59% | 11% |
| PBA_0_ scaffold | 0 | 0 | 0 | n.a. | n.a. |

The values were averages of five scaffolds. Take PBA_1.0_ scaffold as an example, each scaffold was prepared from 20 µL of 1 M PBA precursor solution, having an average dry weight of 13.05 mg. The theoretical adenosine (ADO) sequestration capacity of PBA_1.0_ scaffold is calculated as 5.34 mg if all the feeding PBA molecules were copolymerized and each of them conjugated to an ADO molecule. Thus, the ADO sequestration efficiency ($\eta$) of the PBA_1.0_ scaffold was determined as follows:

$$\eta=\frac{Actual ADO sequestration}{Theoretical ADO sequestration \times PBA incorporation \mathrm{efficiency}}\times100\%$$

$=\frac{3.66 mg}{5.34 mg \times0.915}\times100\%$

$=75\%$

The ADO loading capacity (*%LC*) of PBA_1.0_ scaffold was calculated:

$$\%LC=\frac{Actual ADO sequestration}{Scaffold weight}\times100\%$$

$$=\frac{3.66 mg}{13.05 mg}\times100\%$$

$$=28\%$$

**Figure 1. SEM images of the macroporous scaffolds.**

***a,*** Pore architecture of the PBA_1.0_ scaffold. ***b,*** Pore architecture of the PBA_0_ scaffold.

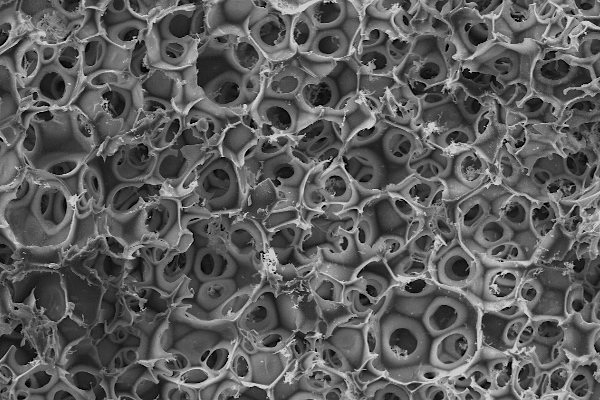

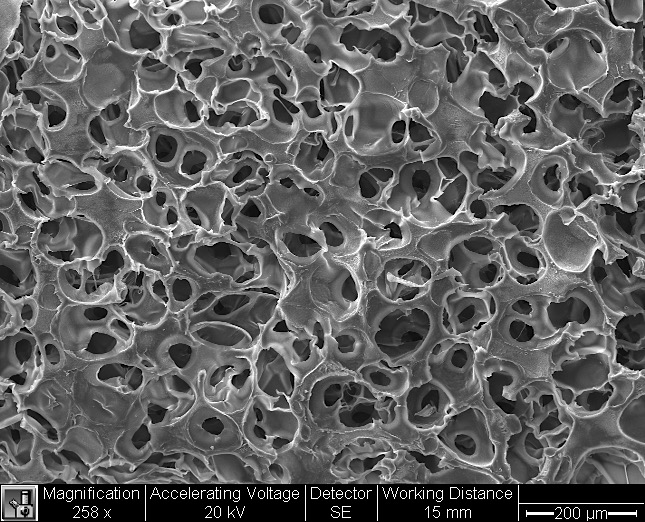

**200 µm**

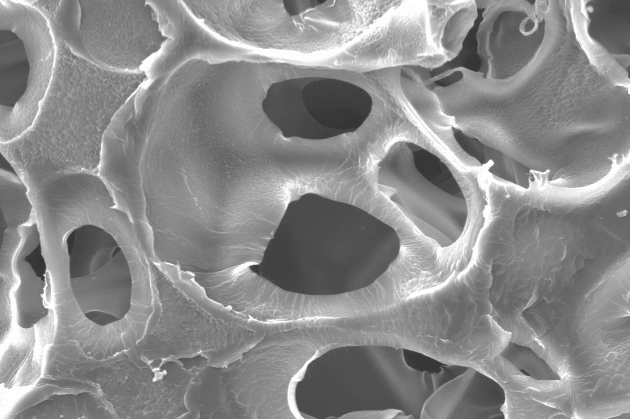

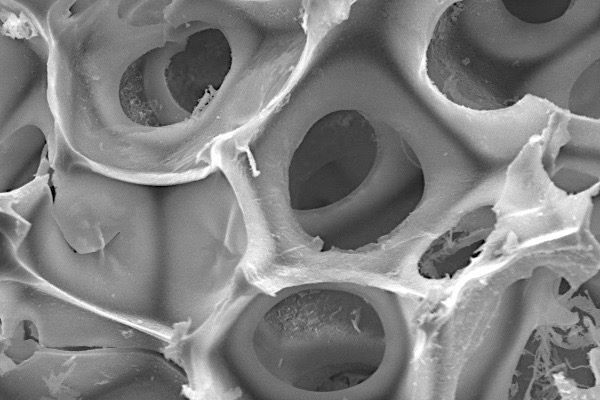

**50 µm**

**a**

**b**

**Figure 2. ^1^H NMR analysis of the scaffolds.**

***a,*** The PEG (PBA_0_) scaffold showed peak at 3.5 ppm, corresponding to −(O*CH_2_CH_2_*)_n_− protons from PEG. Also, peaks from 1.1-1.85 ppm corresponding to −*(CH_2_*)*_4_*− protons indicated the presence of 6-aminocarpoic acid^2^. ***b,c,*** The PEG scaffolds incorporated with PBA (0.5 M and 1 M, respectively) showed an appearance of new peak at around 7 ppm, clearly indicating the presence of aromatic protons and the successful incorporation of PBA into the scaffolds.

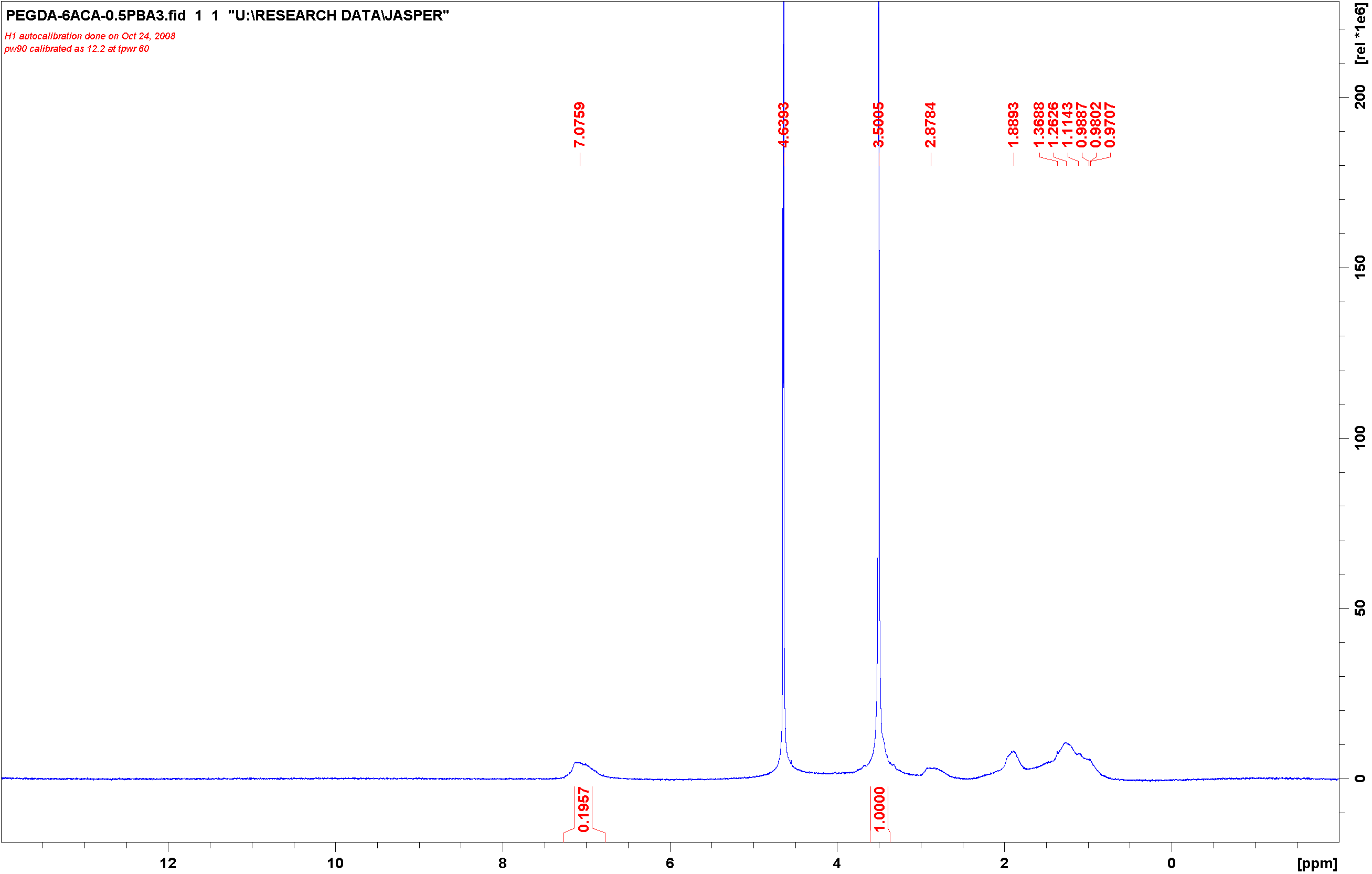

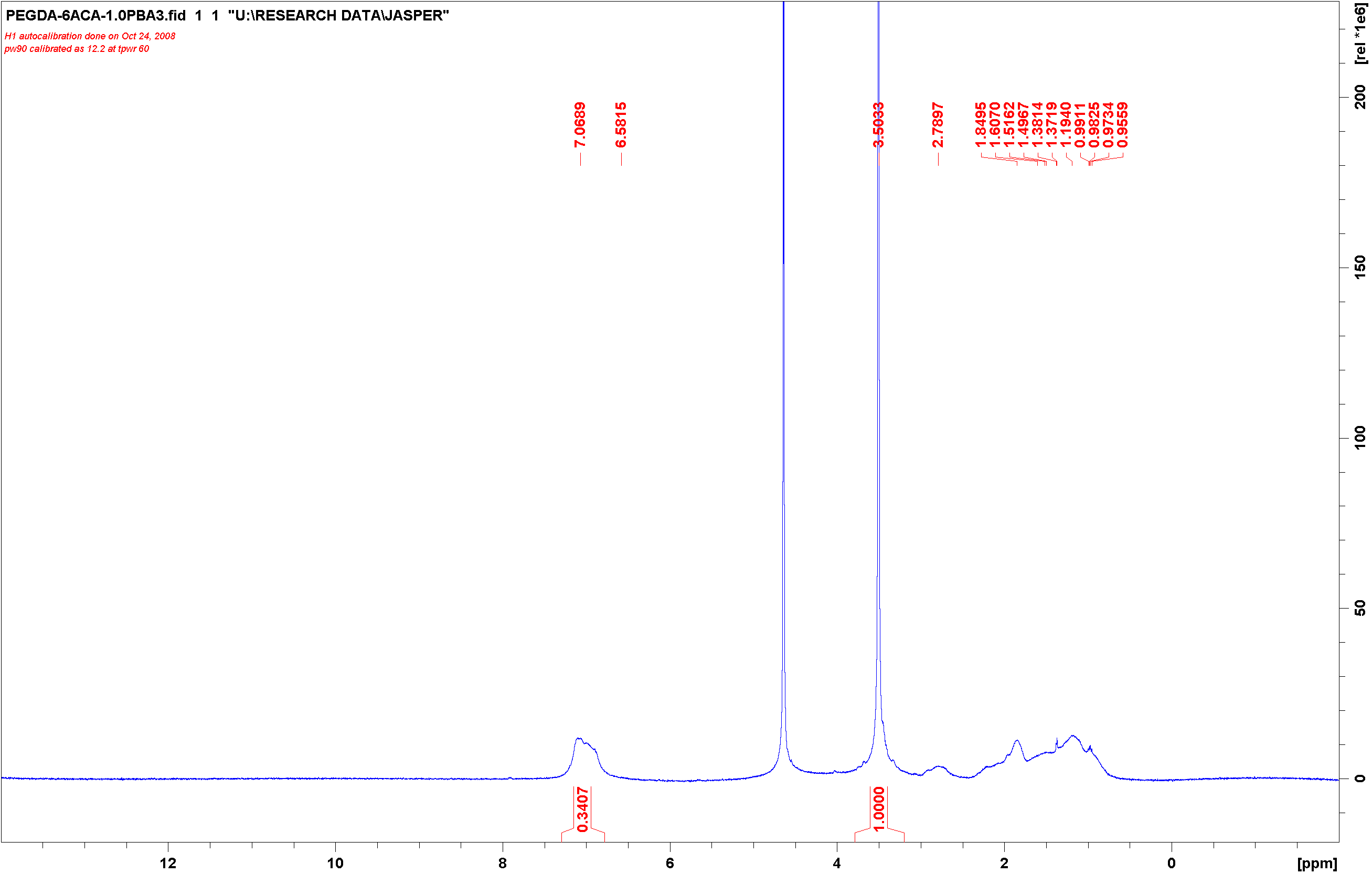

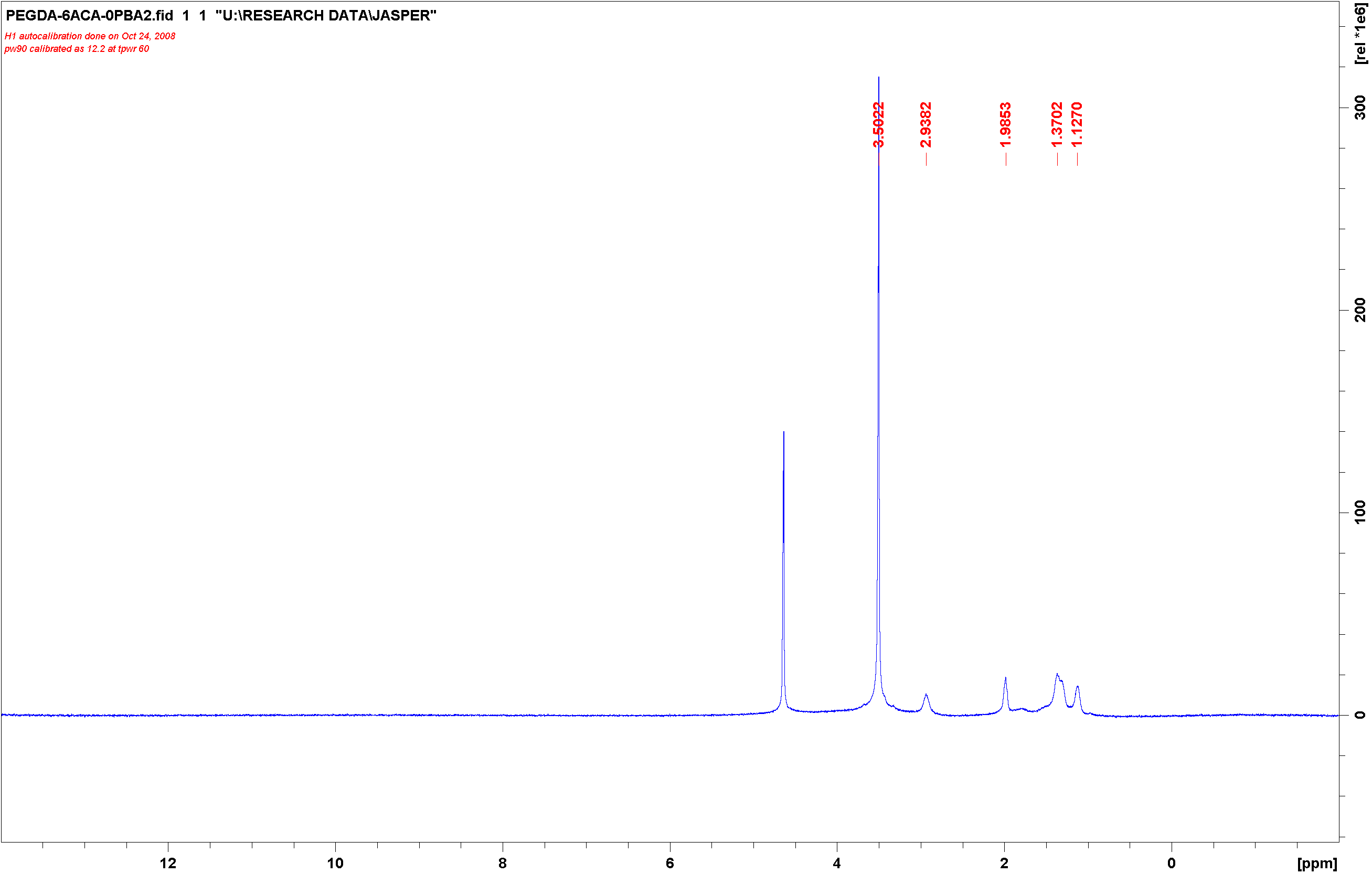

[ppm]

[ppm]

[ppm]

**a**

**b**

**c**

**PBA_0_**

**PBA_0.5_**

**PBA_1.0_**

**Figure 3. Remaining adenosine in the scaffolds at 21 d post fracture.**

Macroporous scaffolds (n = 4) were retrieved at 21 d after fracture and analyzed for their adenosine content using Adenosine Assay. Significance is determined as **p < 0.01 and ***p< 0.001.

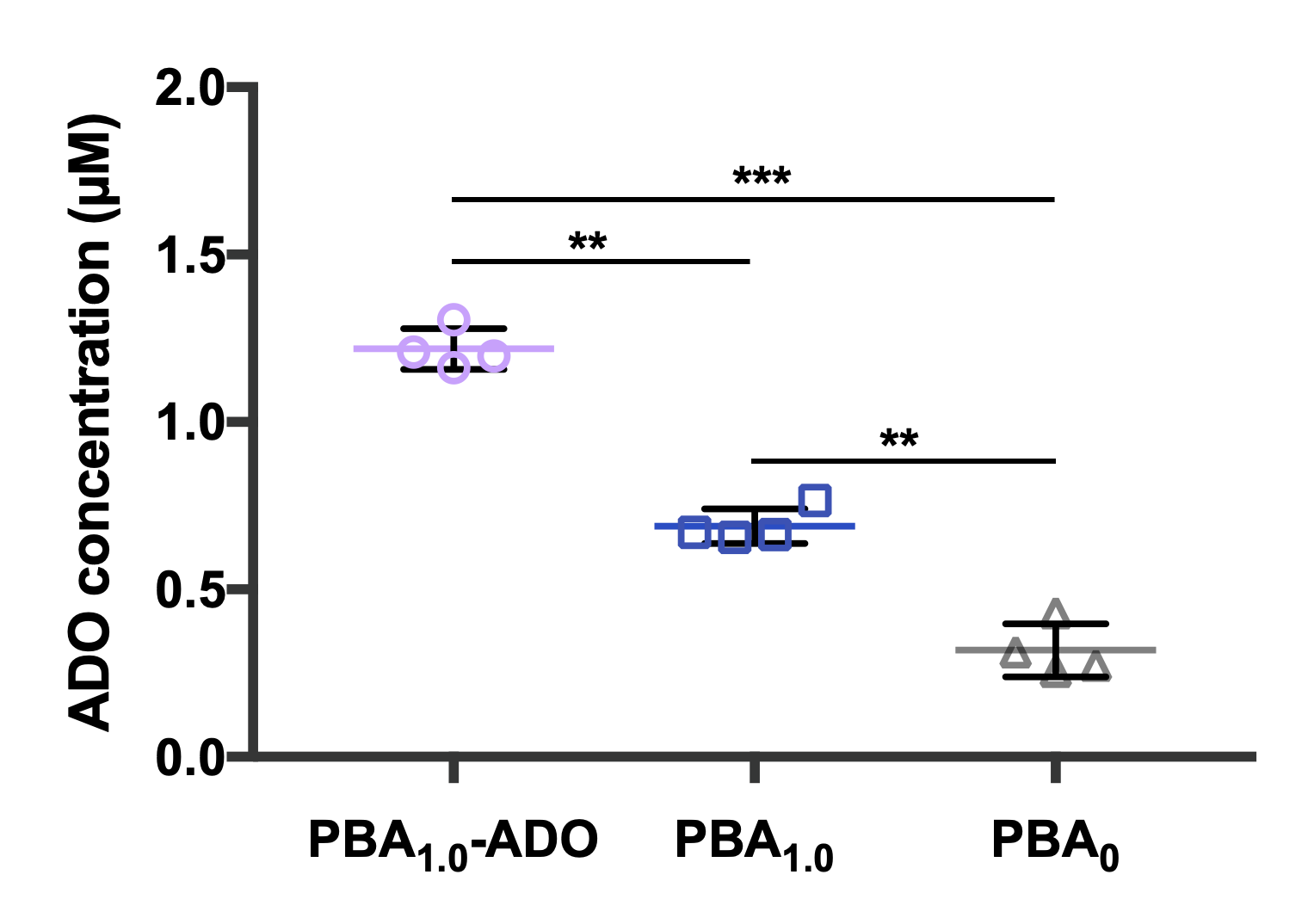

**Figure 4**. **Viability analysis of cells seeded in the scaffolds.**

PBA_1.0_ and PBA_1.0_-ADO scaffolds loaded with hMSCs were cultured *in vitro* in different culture media (GM: growth medium, OM: osteogenic-inducing medium) for 7, 14, and 21 d (n = 3). The DNA content in the scaffolds as a function of time was quantified following a fluorescence method at 480 nm/520 nm (excitation/emission)^3^.

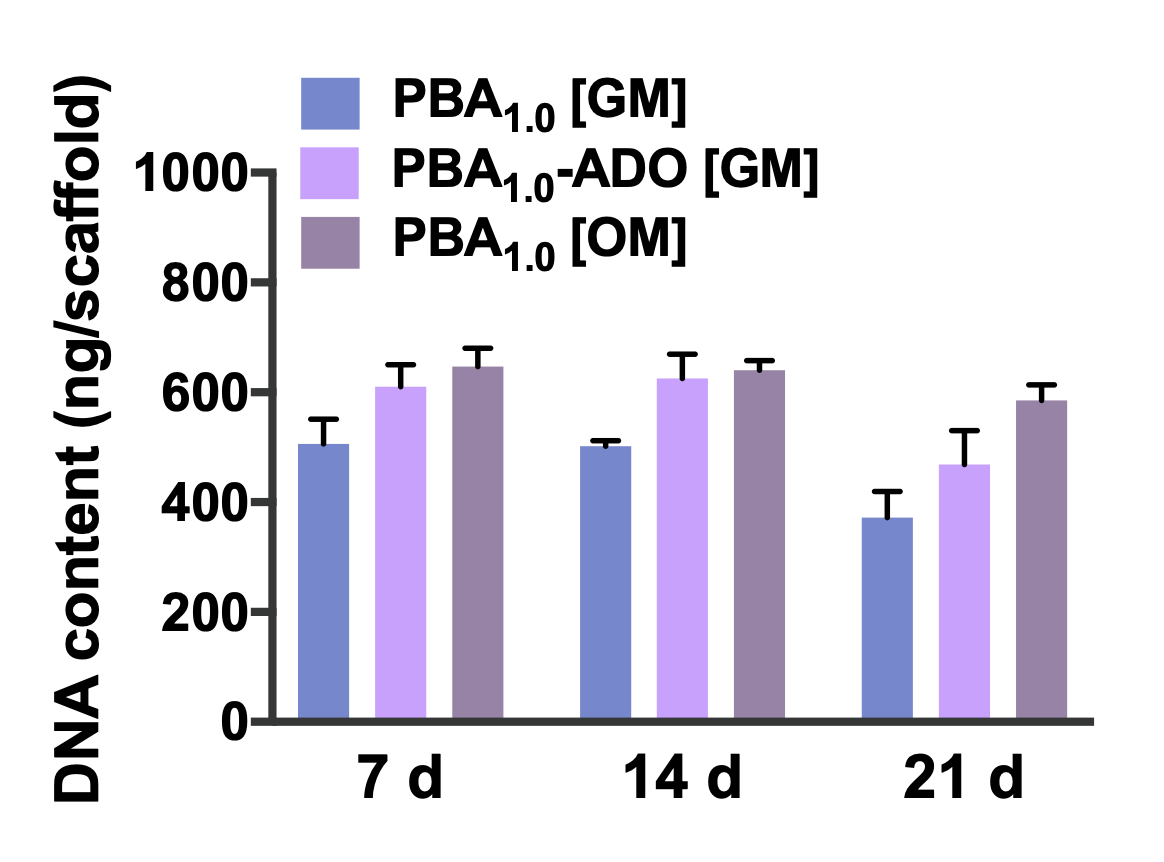

**Figure 5**. **Gross examination of the scaffolds retrieved at 28 d after subcutaneous implantation.**

Both PBA_1.0_ and PBA_1.0_-ADO scaffolds loaded with hMSCs were implanted subcutaneously for 28 d and excised to examine the tissue formation.

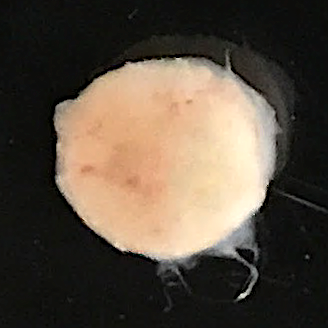

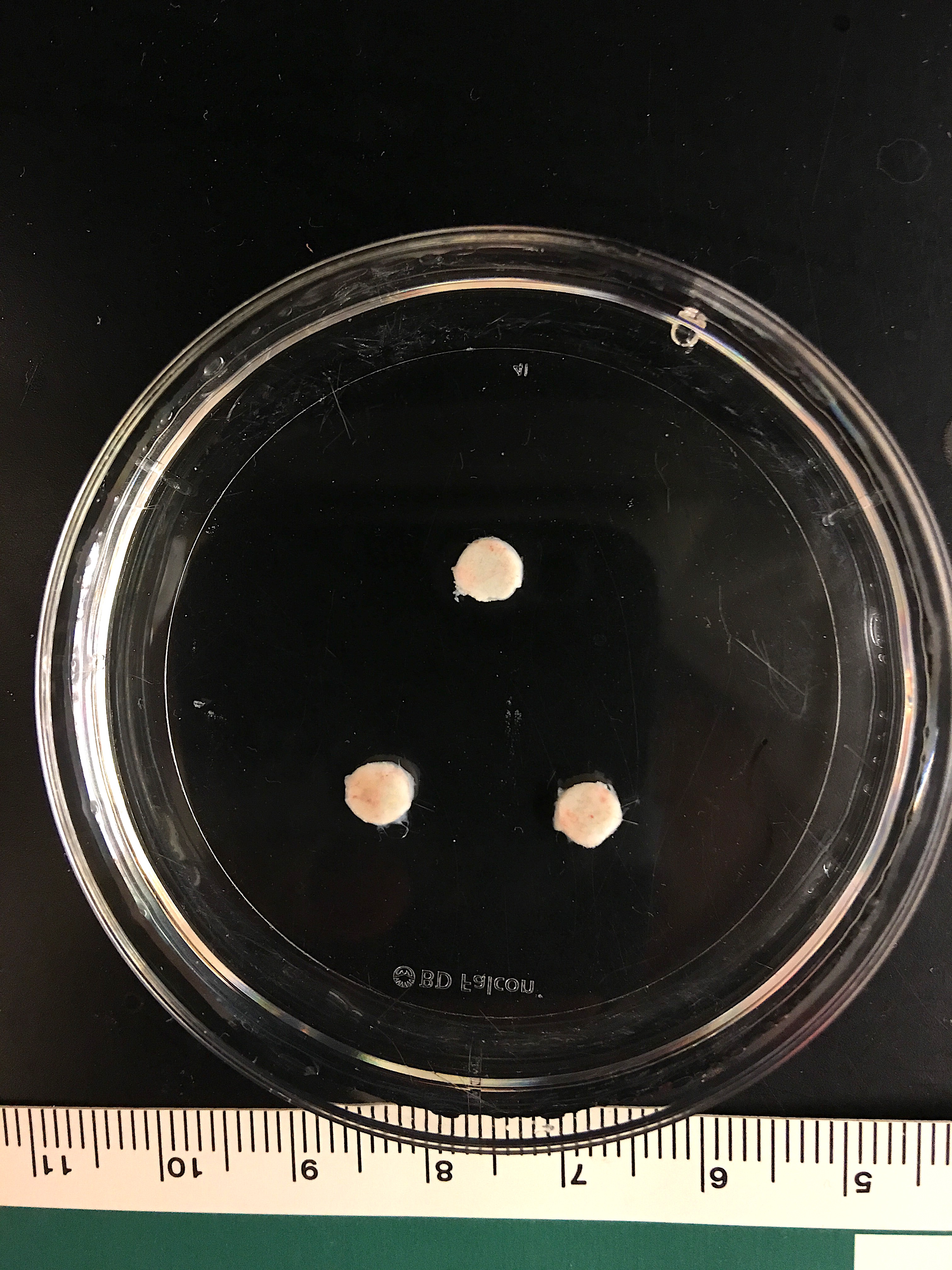

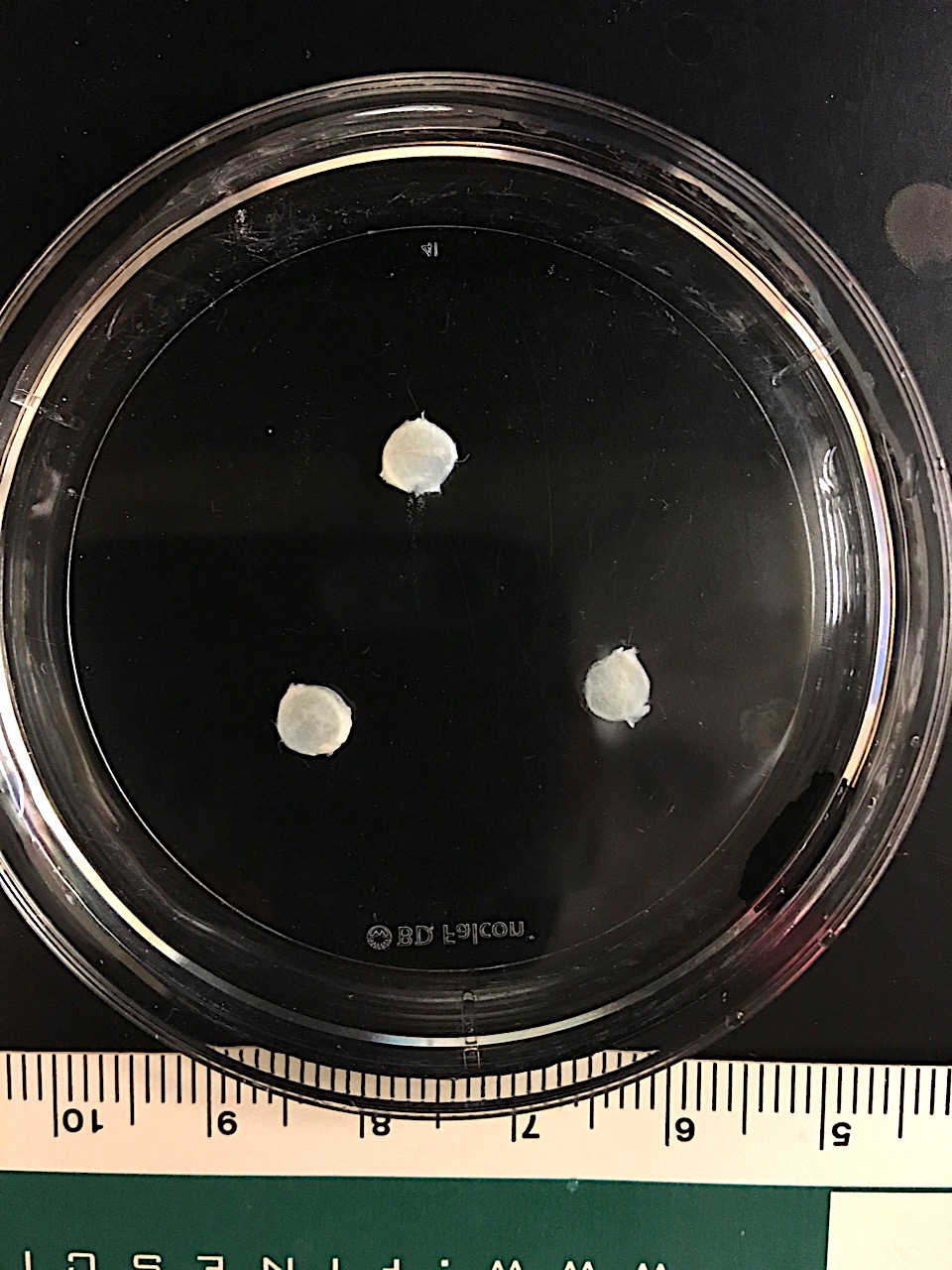

**PBA_1.0_-ADO scaffolds w/ hMSCs**

**PBA_1.0_ scaffolds w/ hMSCs**

**1 mm**

**1 cm**

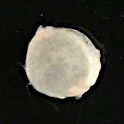

**Figure 6. TRAP** **staining of osteoclasts within calluses.**

***a,*** Tibial fractures were treated with PBA_1.0_-ADO, PBA_1.0_, or PBA_0_ scaffolds for 21 d, and osteoclast activity within the calluses was monitored with time through TRAP staining. Osteoclasts were stained in bright red, with nuclei counterstained in blue. ***b,*** Quantification of the TRAP-positive area to the callus area at 21 d based on the images (n = 5 mice from each treatment; each data point is averaged from 5 images for each mouse). Data are presented as means (± s.d.). One-way ANOVA with Tukey’s multiple-comparisons test was used for statistical analysis in ***b***. Significance is determined as **P* < 0.05 and n.s. (not significant).

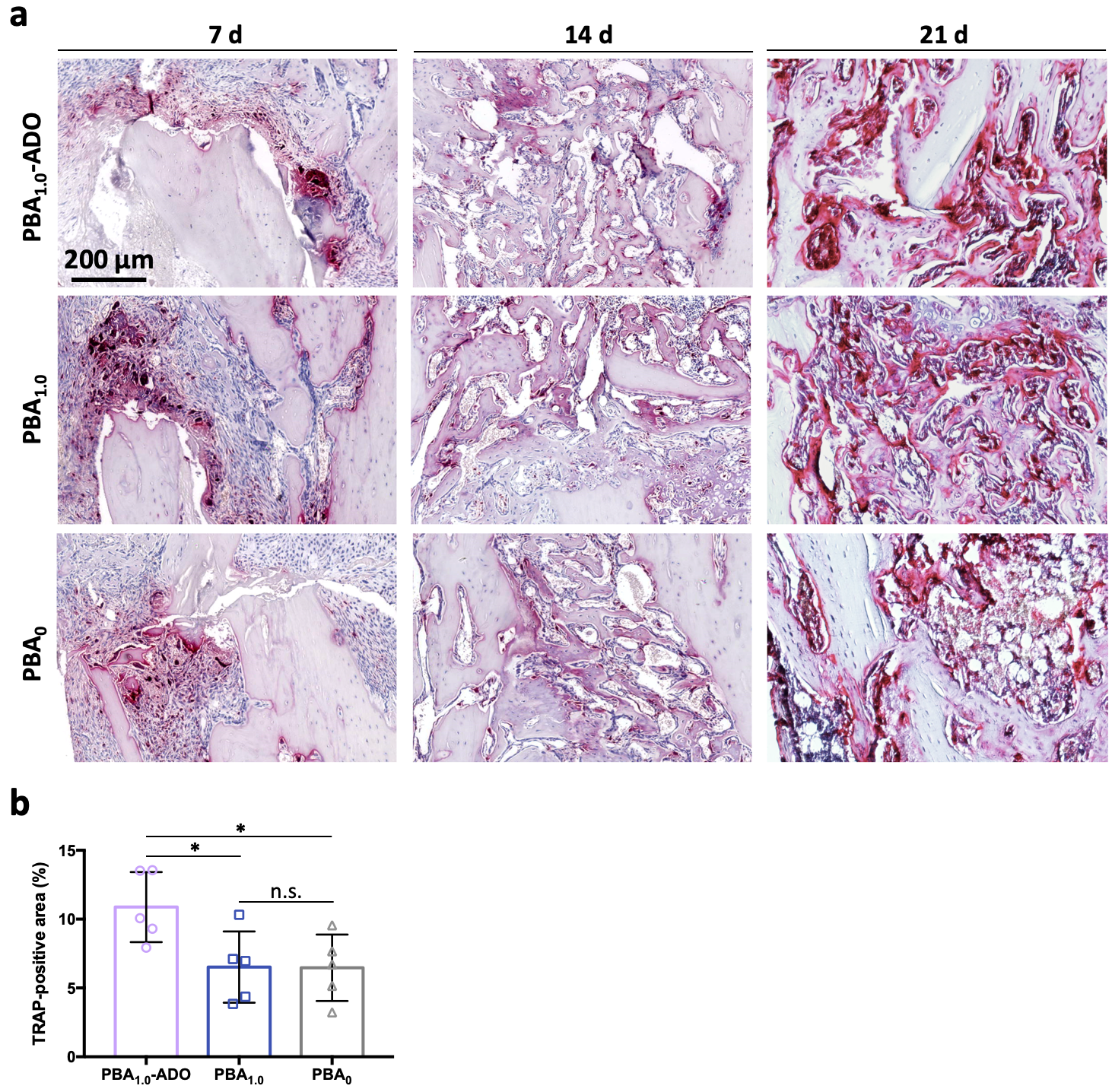
